# Toll-Like Receptor-4 Disruption Suppresses Adipose Tissue Remodeling and Increases Survival During Cancer Cachexia Syndrome

**DOI:** 10.1101/400135

**Authors:** Felipe Henriques, Magno A. Lopes, Felipe O. Franco, Pamela Knobl, Kaltinaitis B. Santos, Luana L. Bueno, Victor A. Correa, Alexander H. Bedard, Adilson Guilherme, Alexander Birbrair, Sidney B. Peres, Stephen R. Farmer, Miguel L. Batista

## Abstract

Cancer-induced cachexia, characterized by systemic inflammation, body weight loss, adipose tissue (AT) remodeling and muscle wasting, is a malignant metabolic syndrome with undefined etiology. Here, we show that Toll-like receptor 4 (TLR4) mediates AT remodeling, in particular, AT browning and inflammatory response in mice bearing Lewis lung carcinoma (LLC). LLC tumor-bearing (TB) TLR4^−/−^ mice were spared from AT remodeling due to a reduced macrophage infiltration and adipocyte atrophy. TLR4^−/−^ mice were also resistant to cold-induced browning of subcutaneous AT (scAT). Importantly, pharmacological inhibition of TLR4 reproduced the main protective effect against AT remodeling found in TLR4^−/−^ TB mice. Moreover, the treatment was effective in prolonging the survival and attenuating tumor mass growth when compared to non-treated-TB animals. Further, tumor-induced elevation of circulating pro-inflammatory cytokines was similarly abolished in both genetic ablation and pharmacological inhibition of TLR4. These data suggest that TLR4 is a critical mediator and a promising therapeutic target for cancer-induced AT remodeling.

**HIGHLIGHTS:**

- Genetic ablation and pharmacological inhibition of TLR4 attenuate adipose tissue remodeling during cancer-associated cachexia;
- TLR4 suppression play an essential role in the browning phenotype induced by cachexia;
- Administration of TLR4 drug inhibitor increase survival and reduces tumor mass growth in tumor bearing mice;
- TLR4 pathway is a promising target for cancer-cachexia therapeutic intervention.

## INTRODUCTION

Cancer cachexia syndrome is characterized by systemic inflammation, body weight loss, remodeling of adipose tissue (AT), and skeletal muscle wasting^1^. Consequently, it results in reduced quality of life, decreased survival, and increased complications due to cancer treatment. Cachexia is the main cause of death in approximately 20% to 30% of all patients with cancer^2^. In cancer cachexia patients, impairment of AT lipid metabolism has been demonstrated and longitudinal studies have established that AT loss precedes muscular atrophy^3^. The changes that characterize AT remodeling comprise: an increase in infiltrated inflammatory cells^4,5^, extracellular matrix rearrangement^4,6^ and the appearance of beige cells^7^ in subcutaneous AT (scAT). The physiological role and consequences of cachexia-induced browning are not known. The current consensus is that multiple factors contribute to cancer cachexia, and therapy requires combinational strategies^8^. Interestingly, pharmacological intervention aimed at attenuating the AT remodeling in cachexia have shown satisfactory results, particularly in mitigating tumor growth and increasing animal survival^9^.

A plethora of evidence from both patients and animal studies suggest a compelling link between activation of the inflammatory pathways and development of cancer cachexia^10,11^. In fact, systemic inflammation has been proposed as a critical feature of cancer cachexia, and anti-inflammatory strategies are considered central to the therapy^10^. It is likely that pathways responding to similar pro-inflammatory cytokines that mediate both sterile and infectious inflammation are critical in cancer cachexia. The Toll-like receptor (TLR) system is potentially one such pathway. In a cancer cachexia animal model, genetic ablation of TLR4 elicited a less severe cachexia with an accompanying lower body weight loss, greater lean body and fat mass, and clinical evidence of reduced wasting compared with the age and weight-matched WT mice^11^. More recently, an elegant study showed that TLR4 is a crucial mediator of cancer-induced muscle wasting due to its integration of catabolic signaling by directly activating muscle protein degradation and indirectly increasing cytokine release^10^. Thus, TLR4 may be a critical therapeutic target for cancer cachexia. The role of the TLR4 pathway in AT remodeling during cancer cachexia development remains unexplored.

Based on these considerations, the present study was designed to further investigate the role of TLR4 on AT remodeling and cachexia development. Using a genetic ablation and pharmacological inhibition of TLR4 we demonstrate that the TLR4 pathway plays an essential role in modulating both thermogenic and pro-inflammatory pathways in fat. Suppression of TLR4 signaling results in a robust resistance to AT remodeling, reduced cachexia, and increased survival. Our study sheds light on TLR4 pathway as a promising target for therapeutic intervention for cachexia.

## RESULTS

### TLR4 deletion attenuates AT remodeling induced by cancer cachexia in TB-mice

During the development of cancer cachexia, AT remodeling arises from morphofunctional and inflammatory changes that result in AT dysfunction. Consistent with these observations, we used the Lewis Lung Carcinoma (LLC) cell line to induce cancer-associated cachexia *in vivo.* After 28 days of cell inoculation, we began to observe the classic cachectic symptoms *in vivo*. WT mice (C57BL/6) showed a reduction in total body weight (12%) after the protocol (Fig. 1A). Adipocyte atrophy in scAT demonstrated by marked reduction in adipocyte size (58.2%) and significantly increased fibrosis demonstrated by total collagen content in the extracellular matrix (Fig. 1B and 1C) were also observed. Furthermore, the presence of CD68 and TNF-α positive staining in scAT was found, while no significant difference in CD3 positive cells in scAT was observed (Fig. 1B). On the other hand, TLR4^−/−^ TB mice did not show signs of loss in the total body weight after the cachexia induction, while also presenting a hindrance in adipocyte atrophy and a reduction in the number of TNF-α and CD-68 positive cells in scAT when compared to WT TB group (Fig. 1A-1C). Taken together, the effects of cachexia on inducing AT remodeling were attenuated in TLR4^−/−^ TB mice.

**Figure 1.**
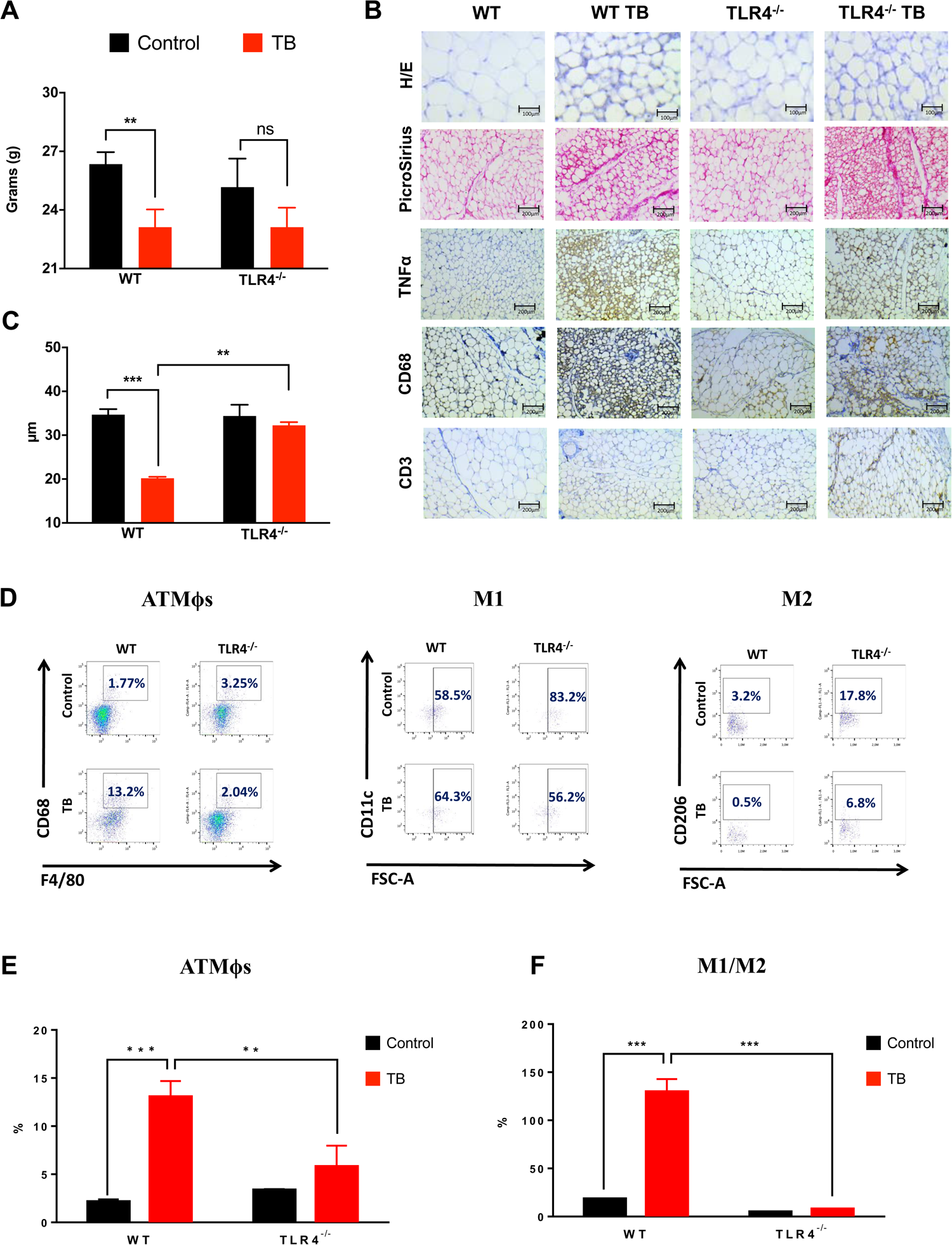
TLR4 deletion attenuates scAT Remodeling during cancer cachexia syndrome. **(A)** Wild-type C57BL/6 and TLR4^−/−^ mice (8-week old male) were inoculated with LLC tumor cells. Total body weight (excluding tumor weight) was evaluated after 28 days before inoculated tumor cells. **(B)** The size of adipocytes (cell diameter) from WT and TLR4^−/−^ mice was quantitatively analyzed (500 adipocytes were measured for each group) after the experimental protocol. **(C)** Histologic sections of scAT in different experimental groups. Histological staining for H/E and picrosirius red were performed, and also, immunohistochemistry for inflammatory profile (TNF*α*) and immune cell markers (CD68 and CD3). **(D)** Stromal vascular fractions (SVF) were isolated from scAT by collagenase digestion for each different group. Flow cytometric analysis of SVF was conducted using fluorescent-conjugated antibodies against CD68, F4/80, CD11c, CD206. Adipose tissue macrophages (ATMϕs) were defined as CD68^+^F4/80^+^ subpopulations and displayed the values as percentage of your respective groups. M1 and M2 ATMϕs were defined as CD68^+^F4/80^+^CD11c^+^CD206^−^ and CD68^+^F4/80^+^CD11c^−^CD206^+^, respectively. These cell populations were shown as a percentage of ATMϕs. **(E)** % of total ATMϕs and **(F)** % of M1/M2 ATMϕs ratio in the scAT. N= 4-5 per group. Scale bars, 100μm and 200μm. Graphs show the mean ± SEM. Statistical significance was determined by Student’s *t*-test or two-way ANOVA. **P < 0.01; ***P < 0.001.

To better understand AT inflammation and the role of macrophage polarization during cancer-associated cachexia, we evaluated AT macrophage (ATMϕs) profiles in scAT from WT and TLR4^−/−^ mice during development of the syndrome. In this case, scAT demonstrated an increase in ATMϕs (6.0-fold) (Fig. 1D and 1E) with an enhancement in M1 (7.0-fold) and a decrease in M2 ATMϕ populations in WT TB mice (Fig. 1D). In this regard, we extended these findings by presenting in greater detail that in models of LLC-induced cachexia, ATMϕs polarization tends to be directed towards an M1 phenotype. On the other hand, the TLR4^−/−^ TB mice presented a consistent attenuation in ATMϕs infiltration (55.3%) and a decrease in the ratio of M1/M2 macrophages (93.2%) in the scAT when compared to WT TB mice (Fig. 1D and 1E). In addition to the macrophage polarization, M2 phenotype are higher in TLR4^−/−^ (4.5-fold) and TLR4^−/−^ TB (12-fold) when we compared with yours respectively controls (Fig. 1D). Therefore, attenuation of TLR4 present a reduction in the recruitment process of ATMϕs thus playing an interesting role during the development of cancer cachexia syndrome.

### Triglyceride turnover is not affected during cachexia in TLR4^−/−^ TB-mice

Considering the attenuation of the AT remodeling seen in TLR4^−/−^ TB mice in response to cachexia, the next step was to evaluate the cachectic response of primary adipocytes (scAT) after isoproterenol (ISO)-stimulus, to evaluate the lipolytic response. During cachexia development, an increase in glycerol released (1.2-fold) in WT TB group was observed relative to the WT group (Fig. 2A). On the contrary, TLR4^−/−^ TB adipocytes showed lower lipolytic values when compared to the TLR4^−/−^ group as well as to the WT TB group (86.8%) (Fig. 2A).

**Figure 2.**
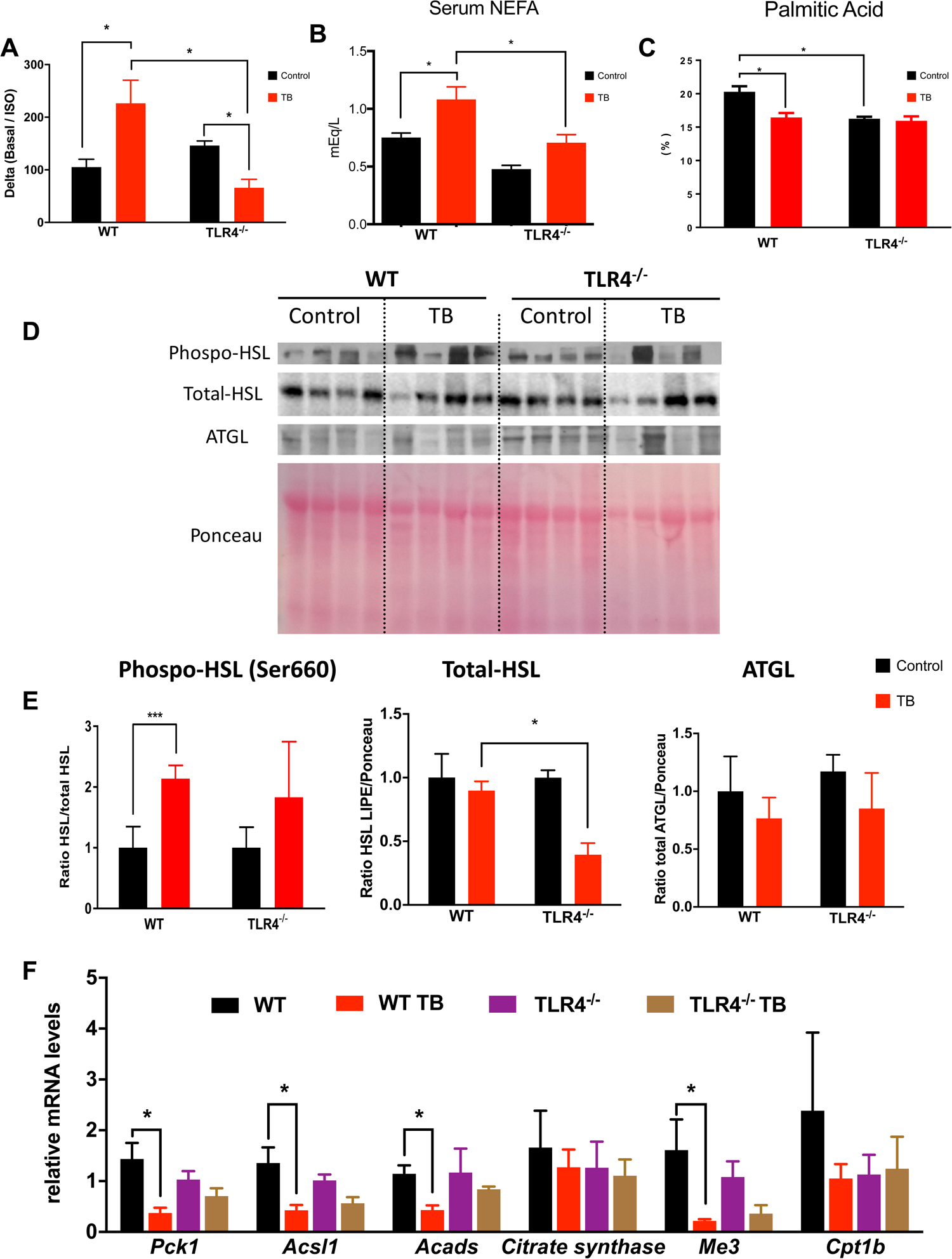
Reduced TG turnover is observed in TLR4^−/−^ TB-mice. **(A)** Primary adipocytes from scAT were incubated with ISO (10μM) for 1 hour to evaluate lipolytic responses. Graph shows the delta (Basal/Iso) glycerol release. **(B)** Serum concentrations of non-esterified fatty acids (NEFA) and **(C)** % of palmitic acid in scAT were performed for each different group. **(D)** Immunoblot analysis for components of lipolysis. scAT lysates were immunoblotted for phospo-HSL (Ser660), HSL and ATGL **(E)** Densitometric evaluation of phospo-HSL (Ser660), HSL and ATGL. Ponceau staining was analyzed as a loading control. **(F)** qRT-PCR was performed to quantitate *Pck1, Acsl1, Acads, Cs, Me3* and *Cpt1b* mRNA levels in scAT from the different groups. N= 4-5 per group. Graph show the mean ± SEM. Statistical significance was determined by Student’s *t*-test or two-way ANOVA. *P < 0.05; **P < 0.01; ***P < 0.001.

Once the attenuation of cachexia-induced lipolysis was shown in TLR4^−/−^ TB mice, serum concentrations of non-esterified fatty acids (NEFA) and palmitic acid levels from the scAT in the different experimental groups were evaluated (Fig. 2B and Fig. 2C). A significant increase was found in NEFA serum concentrations after the cachexia induction in WT TB mice. However, the serum concentration of NEFA was reduced by 36.3% in the TLR4^−/−^ TB mice (Fig. 2B) when compared WT TB. Palmitic acid levels in scAT were reduced by 18.9% in the WT-TB mice when compared with the WT mice (Fig. 2C). There was no cachexic effect in TLR4^−/−^TB (Fig. 2C).

The main lipolytic enzymes were also evaluated. Hormone-sensitive lipase (HSL) showed an increased phosphorylation at ser-660 in WT-TB group, without alteration in the TLR4^−/−^ TB group (Fig. 2D and 2E). There was no change in ATGL level in any experimental condition evaluated (Fig. 2D and 2E). In this scenario, since triacylglycerol (TAG) turnover is affected by cachexia and partially diminished by TLR4 deletion, we evaluated genes involved in fatty acid metabolism in adipocytes. In general, the effect of cachexia on the expression of scAT metabolic markers was reduced in the absence of TLR4. In particular, cachexia reduced expression of *Pck1, Acsl1*, and *Acads* in the WT TB, but not in the TLR^−/−^ TB mice (Fig. 2F).

### TLR4 deletion reduces AT browning and p38MAPK signaling in TB mice

To assess whether TLR4 mediates AT energy expenditure, thermogenic markers were measured in scAT. We found that immunohistochemical staining of UCP1 was increased (1.3-fold) in TB mice (Fig. 3A and 3B). Surprisingly, WT and TB TLR4^−/−^ mice showed a significant reduction in UCP1 staining (Fig. 3A and 3B). Corroborating of this data, the gene profile of the browning “signature” from scAT showed a marked upregulation of *Ucp1* levels in cachectic mice (WT TB), but on the other hand, a significant attenuation in the browning profile was demonstrated in TLR4^−/−^ group, as illustrated in Fig. 3C.

**Figure 3:**
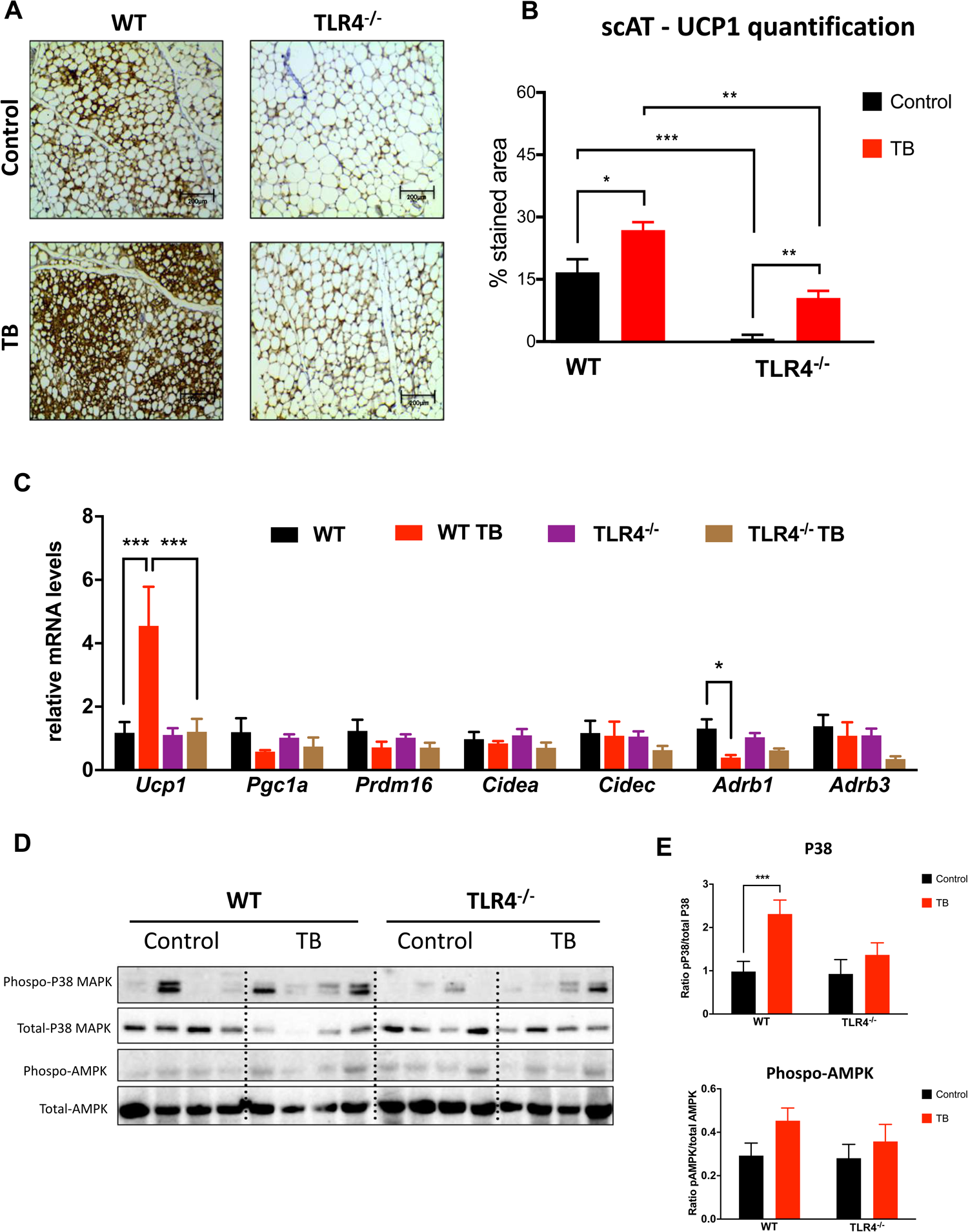
TLR4 deletion reduces browning effect in TB mice throughout p38MAPK pathway. **(A)** Representative images of UCP1 staining of scAT from the different experimental groups. **(B)** Total quantification of UCP1 staining. **(C)** qRT-PCR was performed to quantitate *Ucp1, Pgc1a, Prdm16, Cidea, Cidec, Adrb1* and *Adrb3* mRNA levels in scAT from the different groups. **(D)** Depicted are representative immunoblots to detect phospho-p38MAPK, p38MAPK, phospho-AMPK and AMPK levels in scAT from the different experimental groups. **(E)** Densitometric evaluation of protein levels (phospho/total). N= 4-5 per group. Scale bars, 200μm. Graphs show the mean ± SEM. Statistical significance was determined by Student’s *t*-test or two-way ANOVA. *P < 0.05; **P < 0.01; ***P < 0.001.

Adaptive thermogenesis and UCP1 expression are mainly regulated by sympathetic tone through β-adrenergic signaling and cAMP levels, which can be directly sensed by protein kinase A (PKA) and thus lead to direct or indirect activation of p38 MAPK. Although cachexia induced phosphorylation of p38 MAPK in WT TB, TLR4^−/−^ TB group did not show any change (Fig. 3D and 3E). There were also no changes in the phosphorylation of AMPK (Fig. 3D and 3E) and PKA substrates in any of the experimental conditions evaluated (data not shown) in TLR4^−/−^ TB mice.

### TLR4 is required for cold-induced browning of inguinal white adipose tissue

Having shown that TLR4 is required for browning of scAT during cancer-associated cachexia, we subjected WT and TLR4^−/−^ mice to chronic cold (6°C) challenge to see if TLR4 is required for general adaptive thermogenesis (Fig. 4A). There was a marked reduction in cold-induced browning in TLR4^−/−^ mice compared to WT mice (Fig. 4A and 4B). Interestingly, expression of proteins linked to the lipolytic pathway (HSL, ATGL, and Perilipin) was the same across both groups (WT and TLR4^−/−^), when compared to WT at 22°C (Fig. 4C). Although oxidative metabolism (TCA cycle) was stimulated by chronic cold exposure in both groups (WT and TLR4^−/−^), the increase in *Cpt1b* gene expression was evidenced only in WT, with no change in the TLR4^−/−^ mice (Fig. 4D).

**Figure 4:**
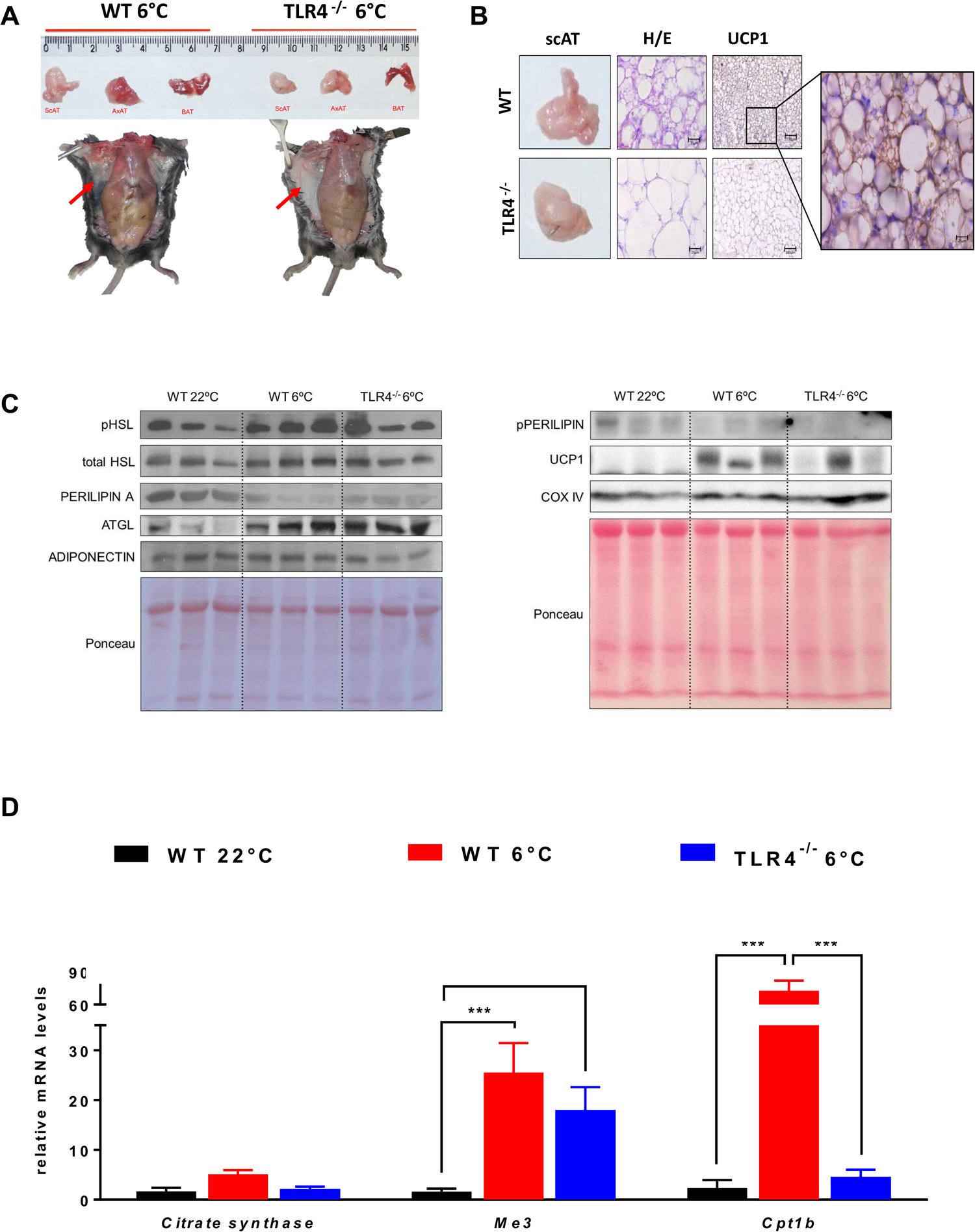
TLR4 is required for cold-inducing browning phenotype. **(A)** Mice were single-caged and housed at room temperature (23°C) and cold exposure (6°C) for six days (n=5 per genotype and per condition). **(B)** Representative images of UCP1 staining in scAT across the different genotypes. Depicted are representative immunoblots to detect phospho-HSL, HSL, perilipin, ATGL, adiponectin, phospo-perilipin, UCP1 and COX IV. **(D)** qRT-PCR was performed to quantitate *Cs, Me3* and *Cpt1b* mRNA in scAT from the different groups. N= 4-5 per group. Scale Bars, 25μm and 100μm. Graph show the mean ± SEM. Statistical significance was determined by Student’s *t*-test or two-way ANOVA. ***P < 0.001.

### Atorvastatin treatment increases survival and attenuates browning

In order to propose a translational approach and to correlate the data with the genetic model, WT TB mice were treated with a 3-hydroxy-3-methyl-glutaryl-CoA (HMG-CoA) reductase enzyme inhibitor, atorvastatin (ATOR). In recent years^12-14^, the anti-inflammatory and pleiotropic effects of this drug have been described, such as inhibition of NFκB activation and downregulation of *Tlr4* and *Myd88* gene expression.

To evaluate if pharmacological attenuation of TLR4 would have an impact on clinical features of cachexia, WT TB mice were treated with ATOR (Fig. 5). The treatment was effective in prolonging the mice survival (Fig. 5A) and also in attenuating scAT atrophy (2.0-fold) and tumor mass growth (49.7%) when compared to TB untreated group (Fig. 5B). Lipolysis was also evaluated in 3T3-L1 cells. Cells that were stimulated with lipopolysaccharide (LPS) and treated with ATOR showed a reduced lipolytic response to the LPS stimulation when compared with untreated cells, showing a significant reduction in 6 hours (41.3%) and 24 hours (45.6%) as illustrated in Fig. 5C.

**Figure 5.**
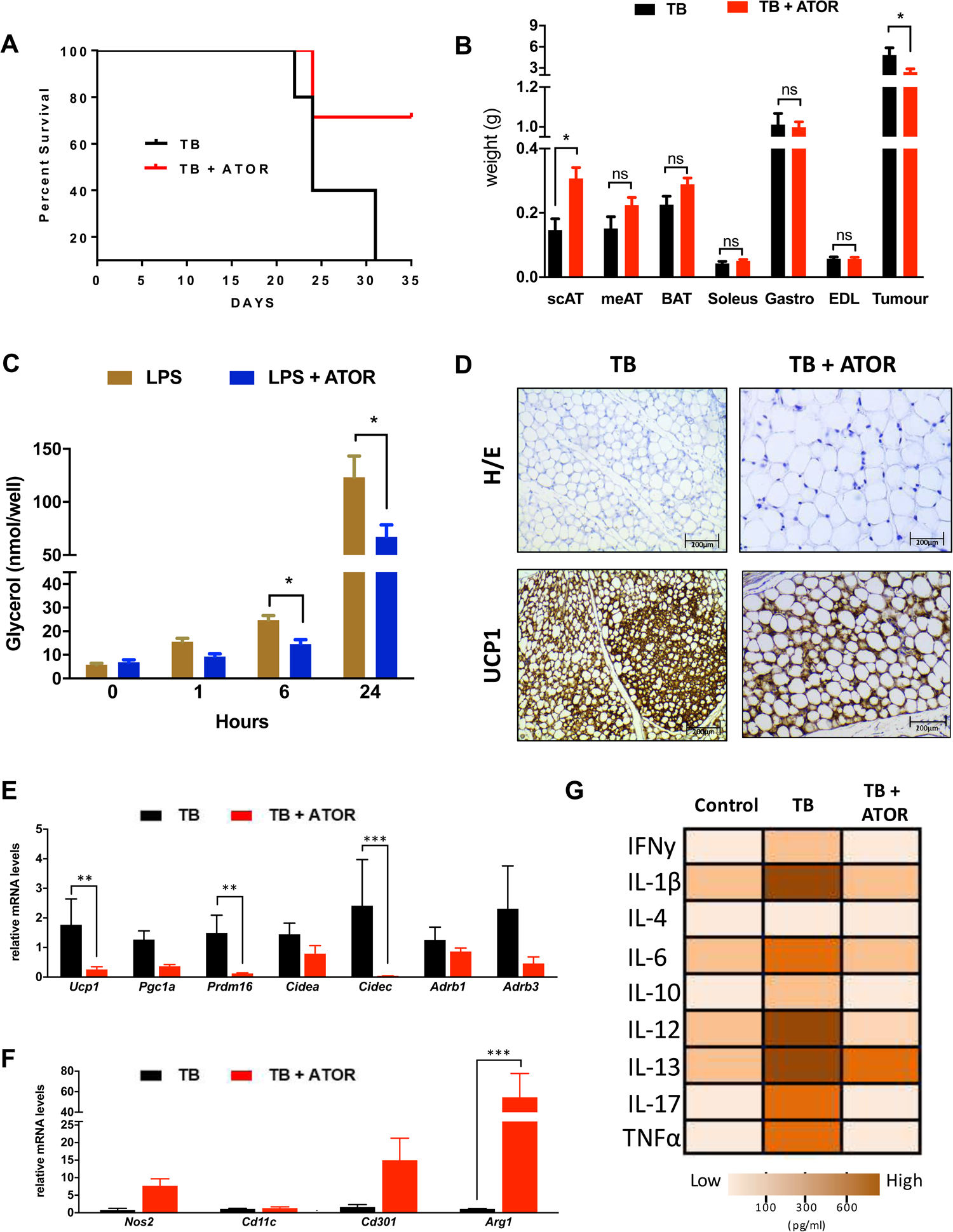
Atorvastatin treatment increases survival and improved cachexia-remodeling in scAT. **(A)** Kaplan-Meier survival curves show a statistically significant difference (P<0.05) in survival between the Tumor Bearing (TB) mice (N=15) and Tumor Bearing + ATOR treatment (TB+ATOR) mice (N=15). **(B)** Adipose tissue, muscle and tumor weights at study end. **(C)** Time course of lipolysis in 3T3-L1 induced by LPS (100ng) and LPS+ATOR treatment (100μM). **(D)** Immunohistochemical analysis for detection of UCP1 in scAT. **(E)** qRT-PCR was performed for *Ucp1, Pgc1a, Prdm16, Cidea, Cidec, Adrb1, Adrb3*, **(F)** *Nos2, Cd11c, Cd301* and *Arg1* for mRNA quantification in scAT from the different groups. **(G)** Heat map representing the circulating pro-inflammatory cytokines in the serum from the different experimental groups. White to brown scale depicts cytokine levels in pg/ml. Data were analyzed using the Bio-Plex manager software. N= 4-5 per group. Scale bars, 200μm. Graph show the mean ± SEM. Statistical significance was determined by Student’s *t*-test or two-way ANOVA. *P < 0.05; **P < 0.01; ***P < 0.001.

Interestingly, ATOR was also able to reduce cachexia-induced browning and atrophy of scAT (Fig. 5D). Furthermore, the profile of the signature browning genes from scAT showed marked downregulation of *Ucp1* (85.5%), *Prdm16* (91.7%) and *Cidec* (98.3%) after ATOR treatment in TB mice, as shown in Fig. 5E. When ATMϕ polarization was analyzed, ATOR treatment appeared to have reversed the previously described M1 ATMϕ polarization within AT.

During cancer cachexia syndrome, systemic inflammation plays a key role in the scAT remodeling, directly influencing the composition of the immune system present in AT, and, consequently, the patient’s immune response. In the present study, we demonstrated that WT TB mice present high plasma concentrations of proinflammatory cytokines in response to cancer-associated cachexia (Fig. 5G). However, after pharmacological treatment with ATOR, it is possible to see a significant decrease in almost all these cytokines, returning the plasma concentration levels to a degree comparable with the WT group (Control group - without cachexia). These results demonstrate that treatment with ATOR is sufficient in improving the inflammatory profile of the cachexic, tumor-bearing mice.

## DISCUSSION

Cancer-associated cachexia is a complex metabolic state accompanied by poor quality of life, high mortality and also resistance to chemotherapy^15-17^. Several clinical interventions to try to improve cachexia through anti-inflammatory drugs and insulin sensitizers but studies have shown limited results^18,19^. Circulating factors produced by the tumor and/or host itself may play an important role for therapeutic treatment for cachexia. Once the factors are identified, they can be targeted for therapeutic intervention.

In our study, we demonstrated that ATOR treatment was effective in increasing the survival of the cachectic mice, in addition to attenuating the main signs and symptoms of cachexia, an effect also observed in the genetic TLR4^−/−^ model. Thus, in addition to the anti-inflammatory and protective effects of skeletal muscle mass as previously demonstrated^10^, we have shown that TLR4 activation is “directly” involved in metabolic disorders in AT such as triglyceride (TG) turnover and browning phenotype, both parameters relevant to the remodeling and dysfunction of AT induced by cachexia. Therefore, the results obtained with the ATOR treatment and TLR4^−/−^ mice during the development of cachexia suggests that the TLR4 pathway plays a fundamental role in maintenance of AT during cachexia syndrome and is a strong candidate for novel anti-cachectic therapy development.

During the development of cancer cachexia, AT remodeling occurs predominantly through morphofunctional rearrangement that is an endpoint process associated with immune-metabolic dysfunction of such tissue^4,5,9,15,23^. In WT TB mice, body weight loss and AT mass wasting were detected, as well as a set of features related to AT remodeling. Interestingly, TLR4^−/−^ TB animal showed to be protected against the effects of cachexia, particularly related to an absence of body weight loss and AT wasting. In obesity (humans and mice model) ^24-26^ a close link between inflammation, production of ECM, development of fibrosis and consequently tissue remodeling was demonstrated in liver and kidney. In these organs, there is mounting evidence that TLR4 acts as a key regulator in fibrogenesis^27-29^ a condition that may or may not be related to TLR4^−/−^ ATMϕs response. However, despite the possible involvement of TLR4 and AT remodeling in cancer cachexia, this topic was not addressed and needs further analysis.

Another relevant aspect of cancer cachexia-induced AT remodeling is the establishment of AT inflammation that was recently characterized by increased recruitment of ATMϕs, including activated M1 and M2 macrophages^5^. In fact, ATMϕ infiltration of scAT has consistently been demonstrated in different experimental models^30^. This scenario has also been demonstrated in patients with cancer and cachexia^4,5^. However, the polarization of ATMϕs in scAT was addressed only recently^5,7^. In this regard, we extended these findings by presenting in greater detail that in cachexia induced by LLC cells, ATMϕs polarization tends to be directed to an M1 phenotype. On the other hand, the TLR4^−/−^ TB presented a consistent attenuation in ATMϕs infiltration in scAT, but in this phenotype, the polarization tended towards an M2 phenotype

The increase in ATMϕs infiltration, largely owing to the recruitment of M1-polarized (or pro-inflammatory) macrophages, and the accompanying increase in crown-like structures are both well-characterized features^31,32^ and a hallmark of AT inflammation during high fat diet (HFD)-induced obesity. However, when obesity is induced in TLR4^−/−^ mice, the increase of AT inflammation was delayed and moderate^24^. In addition, considering that TLR4 is a key initiator of macrophage response and a modulator of systemic insulin sensitivity^33^, the exact mechanism by which TLR4 deficiency affects AT inflammation remains to be explored and would require the generation of an adipose tissue-specific knockout mouse model for TLR4.

In addition to the morphological and inflammatory dysfunction that result in AT remodeling in response to cachexia, TG turnover is the most well-characterized metabolic disorder associated with this condition^4,5,34^, with increased lipolysis being the mechanism best described in the literature^23,35^. In this way, cachectic WT mice led to higher adipocyte lipolysis, which is in line with recent studies demonstrating that AT lipolysis represents a key factor involved in cancer cachexia in both animals^23,35^ and weight-losing human patients^36^. However, adipocytes from TLR4^−/−^ mice were less effected by cachexia and showed very attenuated response to ISO-stimulation.

As lipolysis was strongly decreased in animals lacking TLR4, we attempted to define the main molecular events that lead to cachectic adipocyte metabolism, and so we analyzed the protein expression of key enzymes involved in lipolytic pathways. Although we do observe a higher level of expression or an activating phosphorylation of HSL ser-660, which could explain the greater amount of adipocyte lipolysis observed in cells from cachectic mice (WT TB), such effect was not observed in cachectic TLR4^−/−^ mice. In fact, the increased activity of HSL has been demonstrated in response to cachexia^5,21,37,38^. Even more recently, modulation of lipolysis in different experimental models of cachexia has been shown to be more related to reduced activation of phosphorylation of HSL ser-565 (inhibitory site) and phosphorylation of AMPK, which would increase the lipolytic activity of AT ^21,23^. In this sense, we demonstrated that the reduction in lipolytic capacity observed in the TLR4^−/−^ TB mice was might be followed by changes in phosphorylation of HSL ser-660 induced by cachexia. However, additional studies should address these topics for a better understanding of the pathways involved in modulating lipolysis under these conditions.

Once TLR4^−/−^ TB mice were found to be resistant to the lipolytic effects induced by cachexia, we also evaluated some parameters related to the cycle of NEFA release from the breakdown of stored TGs and re-esterified to TGs (TG turnover). In human cachectic patients, enhanced of TG turnover activity in AT has been shown, as determined by metabolic labeling assay^17^. In our study, WT TB animals showed a reduction of palmitic acid level in the scAT, corroborating the other results regarding the FFA increased by cachexia, a fact that was not present in the TLR4^−/−^ TB mice. Interestingly, this process showed less intensity in TLR4^−/−^ TB mice, including AT morphofunctional preservation. These data substantiate those found in some recent studies that demonstrated an impairment of TG turnover by cachexia^5,35^. In this scenario, since TG turnover is affected by cachexia and partially diminished by TLR4 deletion.

In general, the effect of cachexia on expression of AT metabolic markers was attenuated in the absence of TLR4, particularly in genes related to glyceroneogenesis and TG reesterification. In fact, the role of TLR4 as a physiological regulator of fuel metabolism has been proposed^39^. In this study, during fasting, TLR4^−/−^ mice showed lipid abnormalities that might result from increased TG mobilization, possibly through increased fatty acid reesterification in scAT, decreased in fatty acid oxidation, and/or increased *de novo* lipogenesis in key metabolic tissues. These results corroborate those presented in this study, demonstrating that, in addition to reduced lipolysis, TLR4 deletion may also be involved in the preservation of the deleterious modifications in fatty acid reesterification, induced by cachexia.

It has recently been shown that cachexia induces browning of AT in addition to changes in immune-modulatory activity. In this scenario, chronic inflammation and β-adrenergic activation of thermogenesis functionally cooperate in the pathogenesis of cachexia^3,7^. In the present study, LLC-induced cachexia resulted in an increase in UCP1-positive adipocytes in AT of WT TB mice, followed by a marked upregulation of *Ucp1* levels with absence of upregulation of *Pgc1α, Prdm16, Cidea*, and *Cidec*. These results may be due to the assessed experimental period of 28 days (cachexia staging). Additionally, these results are in line with those presented in other studies with the same experimental model^40^.

Interestingly, TLR4^−/−^ TB showed a significant reduction browning of AT, as well as phosphorylation of p38 MAPK levels, both induced by cachexia. In this sense, the browning phenotype induced by cancer cachexia might be activated indirectly by p38 MAPK and the presence of TLR4 appears to be important in this setting. Recently, it was demonstrated that phosphorylation of p38 MAPK activity was higher in the UCP1 positive axillary AT of cachectic (K5-SOS) animals^7^. Adaptive thermogenesis and UCP1 expression are mainly regulated by sympathetic tone through β-adrenergic signaling and cAMP levels, which can be directly sensed by protein kinase A (PKA) and thus lead to direct or indirect activation of p38 MAPK^41,42^. In this sense, treatment with selective β3-adrenergic receptor (β3-AR) during 4 weeks ameliorates cachexia through attenuation of browning in axillary AT from K5-SOS. However, a well-designed recent study using experimental models demonstrated that full activation of adipose browning, including *Ucp1* expression, in mouse and human cachexia seems to be variable, and the functional contribution to overall energy coats needs to be determined^23^.

A translational perspective on cancer cachexia was evaluated, taking into consideration the results of body mass defense and remodeling of AT after atorvastatin treatment. We observed that suppression of TLR4 significantly attenuates AT remodeling (atrophy and browning) induced by cachexia and also prolongs the survival of the mice that received the atorvastatin treatment (Fig. 5). The reduction of browning induced by cachexia has been demonstrated in different studies, through treatment with cyclooxygenase-2 inhibitor (Sulindac), β3-adrenergic receptor antagonist (SR59230A)^7^ and knockout model for parathyroid hormone-related protein (PTHrP)^43^. However, none of the above-mentioned studies have shown results on survival, which is a key parameter to verify the pre-clinical efficacy of pharmacological intervention.

In cachexia, systemic inflammation is highly dependent on the patient’s immune response. Cytokines are produced by host x tumor integration^44^ and, at least in experimental model, IL-6, TNF-α, and IL-1β are major contributors to the wasting syndrome^6,45^. In addition, systemic inflammation mediators have been used as markers of prognosis of the syndrome, in addition to the presence of cachexia^6,46^. In the present study, the major inflammatory cytokines were abrogated in the same way, both TLR4^−/−^ and pharmacological inhibition of TLR4 in TB mice. In fact, systemic activation of TLR4 increases cytokine synthesis and release from various host cells as an innate immune response^10^. Moreover, a recent study showed the presence of elevated levels of TNF-*α* and IL-6 in animals bearing-LLC tumor, which was attenuated in TLR4^−/−^ mice in the same experimental condition (cachexia)^10^, corroborating the findings of our study.

Most of our findings in this study are based on the data obtained from TLR4 knockout mice. Global TLR4 targeting extends our knowledge of the role of this receptor in AT remodeling caused by cancer cachexia. The TLR4 null mouse model has a significant advantage compared with other knockouts since it prolongs survival from cachexia. The global knockout, however, did not allow us to track TLR4 function at specific developmental stages of cachexia. We also do not know whether some of the phenotypes observed here are due to an adipocyte autonomous, adipocyte non-autonomous or systemic effects. Taken all together, we showed in the present study that TLR4 disruption attenuated adipose tissue remodeling and metabolic dysfunction during the syndrome, thus suggesting a possible a therapeutic target for cancer-induced cachexia

## CONCLUSION

Taken together, our combined data from the TLR4 knockout model and ATOR treatment clearly show that TLR4 plays an essential role in mediating tumor-induced AT remodeling and cachexia development. In this way, disturbing TLR4 signaling systemically may prevent scAT remodeling by abrogating, at least in part, its indirect effects. At this point, both the genetic ablation and the pharmacological inhibition of TLR4 were able to attenuate both the remodeling of the AT (atrophy and inflammation) and the metabolic dysfunction (lipolysis) induced by cachexia (Fig. 6). Also, we have presented, for the first time that the TLR4 receptor may play an essential role in the browning phenotype induced by cachexia. Interestingly, TLR4^−/−^ also showed resistance to browning, suggesting that TLR4 receptor plays an essential role in the morphological alterations and homeostatic energy expenditure. Finally, it is clinically noteworthy that animals treated with a TLR4 inhibitor displayed general attenuation of AT remodeling and systemic inflammation (Fig. 6), presenting an increase in the mice survival, which may have a relevant role as an adjuvant treatment option and a possible therapeutic target for cancer cachexia syndrome.

**Figure 6.**
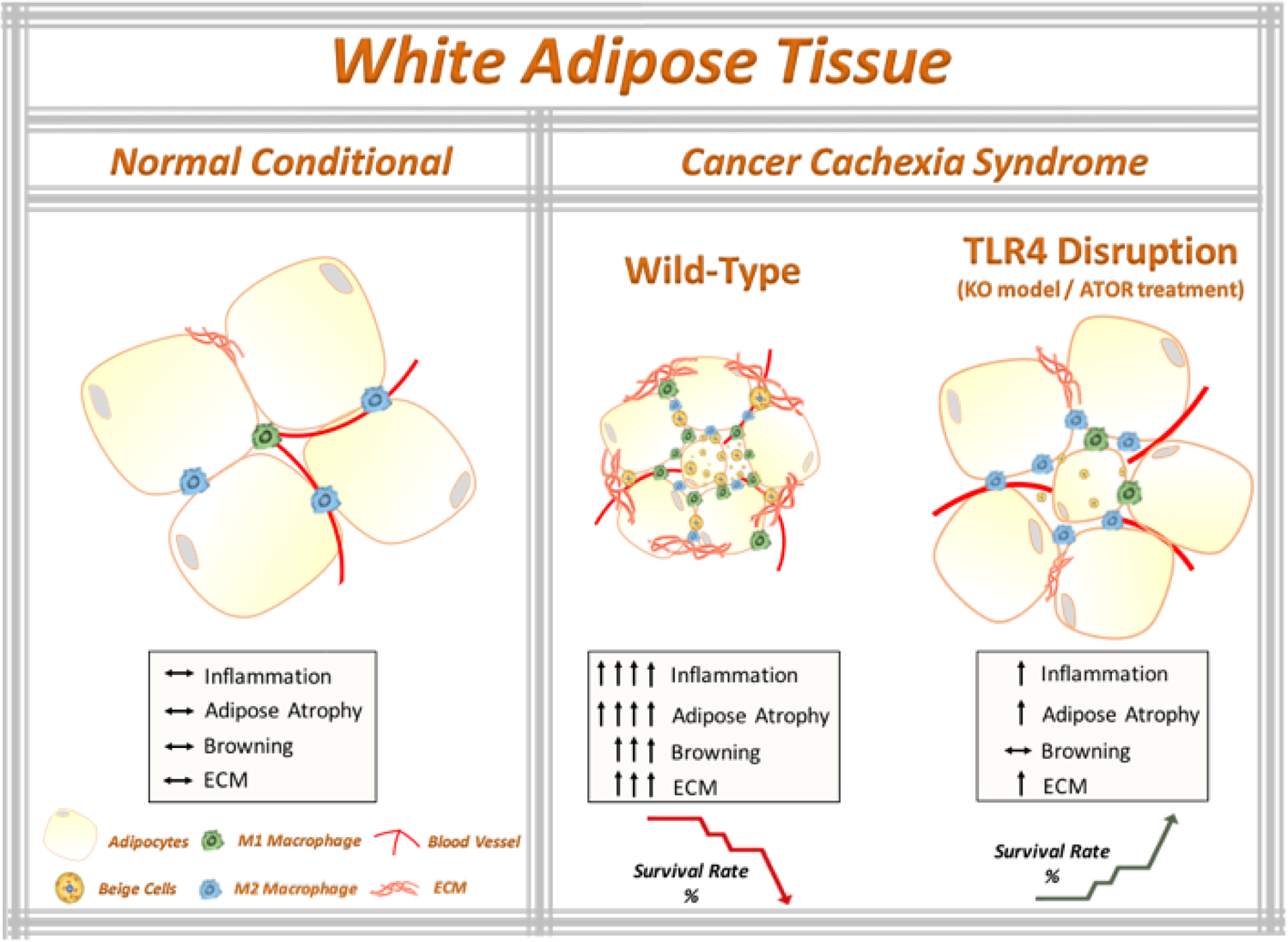
The working model that TLR4 disruption ameliorates adipose tissue remodeling during cancer cachexia syndrome. Cancer-induced cachexia is characterized by systemic inflammation, body weight loss and AT remodeling. We show that TLR4 disruption (knockout model of TLR4 and pharmaceutical inhibition by ATOR treatment) ameliorates AT remodeling, in particular, preservation of adipocyte atrophy and attenuation of browning phenotype in scAT, as well as inflammatory responses during cancer cachexia syndrome. Additionally, TLR4 disruption was effective in prolonging the survival and reducing tumor mass growth. Therefore, these data suggest that TLR4 plays an important role during cachexia development. Here, we suggested a new potential therapeutic target for cancer cachexia syndrome.

## METHODS

### Animal Model

Eight-week old male C57BL/6J and TLR4^−/−^ (Toll like Receptor 4 - Tlr4^lps-del^) mice were obtained from Jackson Laboratory. Mice were housed on a 12 h light/dark schedule and had free access to water and food. Mice were divided into four different groups: Wild Type (WT); WT tumor-bearing (WT TB); TLR4^−/−^ and TLR4^−/−^ tumor-bearing (TLR4^−/−^ TB). For chronic cold challenge, WT and TLR4^−/−^ mice were single-caged and housed at room temperature (23°C) and cold exposure (6°C) for six days (n=5 per genotype and per condition) on a 12h light/12 h dark cycle with access to a standard chow diet. All of the studies performed were approved by the Ethical Committee for Animal Research from the University of Mogi das Cruzes approved all the adopted procedures, which were carried out in accordance with the ethical principles stated by the Brazilian College of Animal Experimentation - Protocol n. 009/2013.

### Tumor Model and Collection of Tissues

Lewis Lung Carcinoma (LLC) tumor cells were used for inducing cancer cachexia syndrome. LLC was injected subcutaneously into the right flanks (200μl LLC cells − 3.5×10^5^). Non-tumor bearing control mice received Phosphate-buffered saline (PBS) only. Development of cachexia was monitored by tumor size and body weight for 28 days. Overnight-fasted mice were euthanized by decapitation without anesthesia, serum was collect and centrifuged (400g, 15min, 4°C) and stored at −80°C for later analysis. After euthanasia, subcutaneous adipose tissue (scAT) was carefully dissected and weighed. Body weight was measured daily, at the same time, over the 28-day experimental period on a precision scale (Ohaus®).

### Atorvastatin Treatment

For the inhibition of the TLR4 receptor via pharmacological action, Atorvastatin (Citalacor®) treatment was perfomed, currently described as a selective antagonist for TLR4. Only WT TB mice were used for this procedure. The animals were divided into two experimental groups: WT Tumor (TB) and WT TB + Atorvastatin (TB+ATOR). Both groups were treated with a concentration of 10 mg/kg/day of atorvastatin by orogastric gavage during the 27-day of protocol. The same protocol was performed for the animals that underwent the survival experiment.

### Histological analysis

scAT was fixed in *HistoChoice® MB (Amresco)*, pH 7.4 for 3h. Fixed tissues were dehydrated and 8-μm sections were used. H&E and Picrosirius staining were performed according to standard procedures (Sigma-Aldrich). The sections were analyzed with a Leica microscope (DM 750). Morphometric aspects were measured by Imagen Pro-Plus 6.0.

### Immunohistochemistry

Immunohistochemistry of scAT was carried out with sections fixed by HistoChoice® MB (Amresco) at pH 7.4 for 3 hours. Fixed tissues were dehydrated and 5-μm sections were used. Dehydrated tissue was blocked using a two different blocking buffers, a buffer solution of endogenous peroxidase activity with 0.3% H_2_O_2_ in methanol and buffer solution of free protein binding sites with 5% normal goat serum. The following primary antibodies were used: UCP1 (ab10983), TNF*α* (ab6671), CD68 (ab955) and CD3 (ab16669). We employed the polymer-peroxidase method, using Histofine Simple Stain MAX-PO (Nichirei Biosciences) and Sigma Fast 3,3-diaminobenzidine as substrate (Sigma-Aldrich). Sections were counter-stained with hematoxylin.

### RNA isolation and qRT-PCR

Total RNA was isolated from scAT using QIAzol Lysis Reagent Protocol (QIAGEN) following the manufacturer’s instructions. cDNA was synthesized from 2 μg of total RNA using iScript cDNA Synthesis Kit (BioRad). Primer sequences used for qRT-PCR analyses were listed in Table S1. Analyses of qRT-PCR products were performed with the Prism 7500 SDS software (Thermo Fisher Scientific). Relative quantification of mRNA amount was obtained by the by 2^−(ΔΔCt)^ method.

### Western Blot

For protein expression analyses, tissues were homogenized in lysis buffer (20 mM HEPES, 150 mM NaCl, 2 mM EDTA, 1% Triton X-100, 0.1% SDS, 10% glycerol, 0.5% sodium deoxycholate) that had been supplemented with Halt protease and phosphatase inhibitors (Thermo Pierce). Samples from tissue lysates were then resolved by SDS-PAGE, and immunoblots were performed using standard protocols. Membranes were blotted with the following antibodies: HSL (ab45422); phospo-HSL Ser660 (Cell Sig. #4126); ATGL (Cell Sig. #2138); P38 MAPK (Cell Sig. #8690); phospo P38 MAPK (Cell Sig. #9211); phospo-AMPK (Thr172-Cell Sig. 2531); AMPK (Cell Sig. 2532); Perilipin (ab3527); phospo-Perilipin (ab87129); Adiponectin (ab62551); UCP-1 (ab10983) and COX IV (ab16056), all overnight. The membranes were incubated with the goat anti-rabbit IgG conjugated to horseradish peroxidase secondary antibody (7074 − Cell Signaling Technology) for 1 h at room temperature and were detected by ECL Prime (Amersham). For the control of protein loading and transfer, we used Ponceau staining of the membranes. Quantification of the antigen–antibody complex was performed by the ImageJ Analysis Software.

### SVF isolation and Flow cytometry analysis

Stroma vascular fraction (SVF) from scAT were excised and minced in PBS with calcium chloride and 0.5% BSA. Collagenase II (Sigma-Aldrich) was added to 1 mg/ml and incubated at 37°C for 30 minutes with shaking. The cell suspension was filtered through a 100 μm filter and then spin down at 300 g for 5 minutes to separate floating adipocytes from the SVF pellet. Cells were incubated with Fc Block (BD Biosciences) prior to staining with conjugated antibodies for 15 minutes at 4°C. SVF were purified and stained with the following antibody panel: ATMϕs were defined as CD68^+^F4/80^+^ subpopulations. M1 and M2 ATMϕs were defined as CD68^+^F4/80^+^CD11c^+^CD206^−^ and CD68^+^F4/80^+^CD11c^−^CD206^+^, respectively. Cells were analyzed using a BD Accuri (BD biosciences). The data was analysis was performed using FlowJo.

### Measurement of plasma cytokines

Plasma was collected using plasma separators (BD Biosciences). Cytokine levels were determined using a Milliplex assay kit (Milliplex MAP mouse cytokine/chemokine magnetic bead panel, #MCYTOMAG-70K - Millipore). Data were analyzed using the Bio-Plex manager software (Bio-Rad Laboratories).

### Lipolysis assay

Isolated adipocytes were suspended in 40 ml of Kreb’s/ Ringer/phosphate buffer (pH 7.4, with 1% BSA) in 0.6 ml microtubes with 20 ml of adenosine deaminase (0.2 U/ml) and were incubated at 37°C for 5 min (as to degrade endogenously released adenosine). Afterward, the cells were incubated for 1 h at 37°C either in presence of isoproterenol (β-adrenergic agonist) for determination of glycerol concentration. For additional analyses of the possible effect of ATOR on inhibiting lipolysis, 3T3-L1 cells were stimulated during 1h, 6h and 24h with lipopolysaccharide (LPS, 100 ng/mL). The concentration in the medium of released glycerol was quantified with a commercially available reagent (Sigma-Aldrich), measured according to the manufacturer’s instructions.

### Cell culture

Swiss preadipocytes 3T3-L1 cell line were plated at 1×10^4^ in 24-well culture plates and cultured in DMEM (Dulbecco’s Modified Eagle’s Medium) with 4500 mg glucose supplemented with 10% bovine serum and 2% penicillin with streptomycin at pH 7.4. The cells were maintained at 37°C with 5% carbon dioxide (CO_2_) so as not to reach complete confluence until they were induced to differentiate. Preadipocytes were brought to complete confluence (day-2) and after two days of confluence, the culture medium was replaced by differentiation inducer medium, consisting of DMEM, supplemented with 10% fetal bovine serum (FBS), 1 μM dexamethasone, 0.5 mM 3-isobutyl-1-methylxanthine (IBMX) and 1.67 μM bovine insulin. From the second day of differentiation, the cells were maintained in culture medium containing only 0.83 μM insulin and 10% FBS which was changed every 48 hours for eight days.

### Statistical analysis

Data were analyzed in GraphPad Prism 7 (GraphPad Software). The statistical significance of the differences in the means of experimental groups was determined by Student’s t-test or and 2-way ANOVA, followed by Tukey’s post hoc comparison tests. The data are presented as means ±SEM. P values ≤ 0.05 were considered significant.

## ACKNOWLEDGMENTS

We thank all members of the Laboratory of Adipose Tissue Biology for helpful discussions and critical reading of the manuscript. This work was supported by São Paulo Research Foundation (FAPESP) Grants: 2010/51078-1, 2015/19259-0 and CNPq 311966/2015-2 to M.L.B.Jr. The contents of this article are solely the responsibility of the authors and do not necessarily represent the official views of FAPESP.

## AUTHOR CONTRIBUTIONS

F.H, M.L.B.Jr, S.B.P and S.R.F conceived the study; F.H, M.L.A, F.O.F, P.K, L.L.B,

A.H.B and K.B.S performed the research; A.G, A.B, S.B.P, S.R.F, F.S.H, M.L.A,

F.O.F and M.L.B.Jr analyzed the data; F.H and M.L.B.Jr wrote the paper.

